# Binocular disparity-based learning does not generalize to the other visual hemifield

**DOI:** 10.1101/2020.08.17.250985

**Authors:** Jens G. Klinzing, Hendrikje Nienborg, Karsten Rauss

**Author notes:** Authors for correspondence: Jens G. Klinzing and Karsten Rauss Otfried-Mueller-Str. 25, 72076 Tübingen, Germany.

## Abstract

Visual perceptual learning refers to long-lasting performance improvements on a visual skill, an ability presumably supported by plastic changes in early visual brain areas. Visual perceptual learning has been shown to be induced by training and to benefit from consolidation during sleep, presumably via the reactivation of learning-associated neuronal firing patterns. However, previous studies have almost exclusively relied on a single paradigm, the texture discrimination task, on which performance improvements may not rely on strictly perceptual skills. Here, we tested whether sleep has beneficial effects on a visual disparity discrimination task. We confirm previous findings in showing that the ability to discriminate different disparities is unaffected by sleep during a 12-hour retention period after training. Importantly, we extend these results by providing evidence against an effect of sleep on the generalization of improved disparity discrimination across the vertical meridian. We further rule out carry-over effects as a possible confound in our earlier within-subject study by relying on a between-subject design. The combined data from both studies argue against sleep as an important factor in the consolidation of a strictly perceptual skill. This sets important constraints on models of the role of sleep and sleep-associated neural reactivation in the consolidation of non-declarative memories.

## Introduction

In biological memory systems, recently acquired information is often susceptible to loss and distortion. A subset of novel memories thus undergoes an active process of consolidation^1,2^. This process, often referred to as ‘systems consolidation’, is critically supported by repeated reactivations of neuronal activity patterns representing the acquired information during post-learning sleep^3^. Successful consolidation reduces the probability of information to be forgotten, but it also induces qualitative changes to the information, often including improved generalization. For visual learning, this may encompass a higher invariance to specific properties of a stimulus, such as its location in the visual field.

Visual learning is commonly investigated using the classical texture discrimination task (TDT). Improvements on this task tend to be confined to the trained eye or specific regions of the visual field. This suggests that visual learning is supported by synaptic changes in early stages of visual processing, where information is still retinotopically organized^4^. This notion is supported by higher Blood-oxygen-level-dependent imaging responses in the visual cortex to the TDT when stimuli were presented to the trained compared to the untrained eye^5^. A series of studies showed benefits of sleep on improvements in the TDT^6–13^, including the ability to generalize to the untrained eye^12^. Due to the retinotopic organization of early visual areas, local mechanisms are unlikely to account for this effect, suggesting a role of top-down feedback. Coordinated memory reactivation between hippocampus and visual cortex, potentially reflecting such top-down mechanism, has been shown in sleeping rats^14^. Other studies show an essential role of the thalamus in consolidating plastic changes in the visual cortex^15,16^. Together these findings suggest newly acquired visual skills may benefit from sleep via interactions of the visual system with higher-order areas during sleep.

As a major caveat to this idea, sleep benefits on visual learning have almost exclusively been demonstrated on the TDT. Importantly, difficulty in the TDT is adjusted by changing the time window between the target stimulus and a subsequent mask, not by changing properties of the target stimulus itself. Improvements on the task may therefore result from better temporal allocation of attentional resources. In line with this idea, training subjects on temporal aspects of the task beforehand has been shown to minimize further improvements on the TDT^17^. One study showed that sleep may benefit improvements on the extraction of a prototype from abstract visual shapes^18^. Higher performance on this task may thus result from improvements in higher-order processing and not on its perceptual component.

In a previous study^19^, we therefore investigated the effect of sleep on a strictly perceptual task. This task required coarse binocular disparity discrimination, the ability to use disparities in the image received by both eyes to assess depth. In primates, first signs for the processing of binocular disparity are found in the primary visual cortex^20^, such that overnight improvements in the task are likely to result from top-down feedback. We investigated potential benefits of sleep on binocular disparity discrimination in terms of improvements at the trained location as well as generalization across the horizontal meridian of the visual field. In that previous study, we showed that subjects successfully improve performance in the trained task, however, this effect was specific for the trained location. We did not demonstrate generalization of the learned skill from the lower to the upper visual field. This effect was independent of whether the time between learning and testing was spent awake or asleep. A possible confounding factor concerning these previous results are potential differences in baseline performance between the lower and upper hemifield. Previous research has shown that in primates, visual processing differs between the lower and upper visual field^21,22^. From an evolutionary perspective, it is plausible that detailed depth discrimination would be more strongly selected for in the lower visual field. Thus, poor baseline performance in the upper visual field could have masked generalization in our previous study. To rule out such a confound, we reproduced our previous study with the important difference that we tested generalization across the vertical instead of the horizontal meridian. Furthermore, we addressed potential carry-over effects between wake and sleep conditions by testing this factor across groups instead of within subjects. In line with our previous study, we present evidence against generalization of binocular disparity learning across the visual meridian. Our results do not rule out the existence of local generalization to close-by receptive fields. However, it is unlikely that early perceptual learning of the kind investigated in this study is subject to global generalization by broad top-down feedback as presumed by systems consolidation.

## Methods

### Participants

32 participants (17 female, 15 male; mean age ± SD: 23.66 ± 3.25, range: 18-30 years) were trained on a binocular disparity discrimination task used previously^19^. Nine additional subjects started the experiment but where excluded because they were not able to see the depth effect and therefore performed at around chance level (n = 7), slept during the retention interval despite being assigned to the wake group (n = 1), or as a result of technical difficulties (n = 1). Subjects were required to score above 1.0 in a decimal visual acuity test, corresponding to a Snellen acuity of 20/20. Exclusion criteria were health problems, ongoing medication, medical interventions, night or shift work, exam periods, stress-intense occupations during three weeks prior to the experiment, or a history of psychiatric, neurological, or sleep disorders. On experimental days, extensive physical exercise, daytime naps, as well as the consumption of alcohol, caffeine, or illegal drugs were prohibited. The study was approved by the local ethics committee of the medical faculty of the University Tübingen. All subjects gave their written informed consent.

### Binocular disparity discrimination task

Participants performed a two-choice disparity discrimination task in the lower visual field (Figure 1A). They discriminated whether a central disk (3° diameter) was protruding (“near”) or receding (“far”) relative to a ring (1° wide) surrounding the disk. Trials were performed at different difficulty levels, varied by the disk’s signal strength (see below). Between trials, only a fixation cross was shown at the center of the screen. After initiating a trial with a button press, the stimulus (disk and ring) was shown for 1.5 sec with a horizontal (left or right depending on condition) and vertical offset (lower visual field) to the fixation cross of 3°. Stimulus and fixation cross then disappeared and were replaced for a maximum of 2 s by two choice targets (small versions of a near and a far stimulus at full signal strength; 2.2° disk diameter; 0.8° ring width) above and below the previous location of the fixation cross. Participants were asked to pick the stimulus they had just seen using an up or down button press (match-to-sample). From stimulus to choice target onset, eye fixation was assessed using an infrared eye tracker (Eyelink 1000, SR Research) at a sampling rate of 500 Hz.

**Figure 1.**
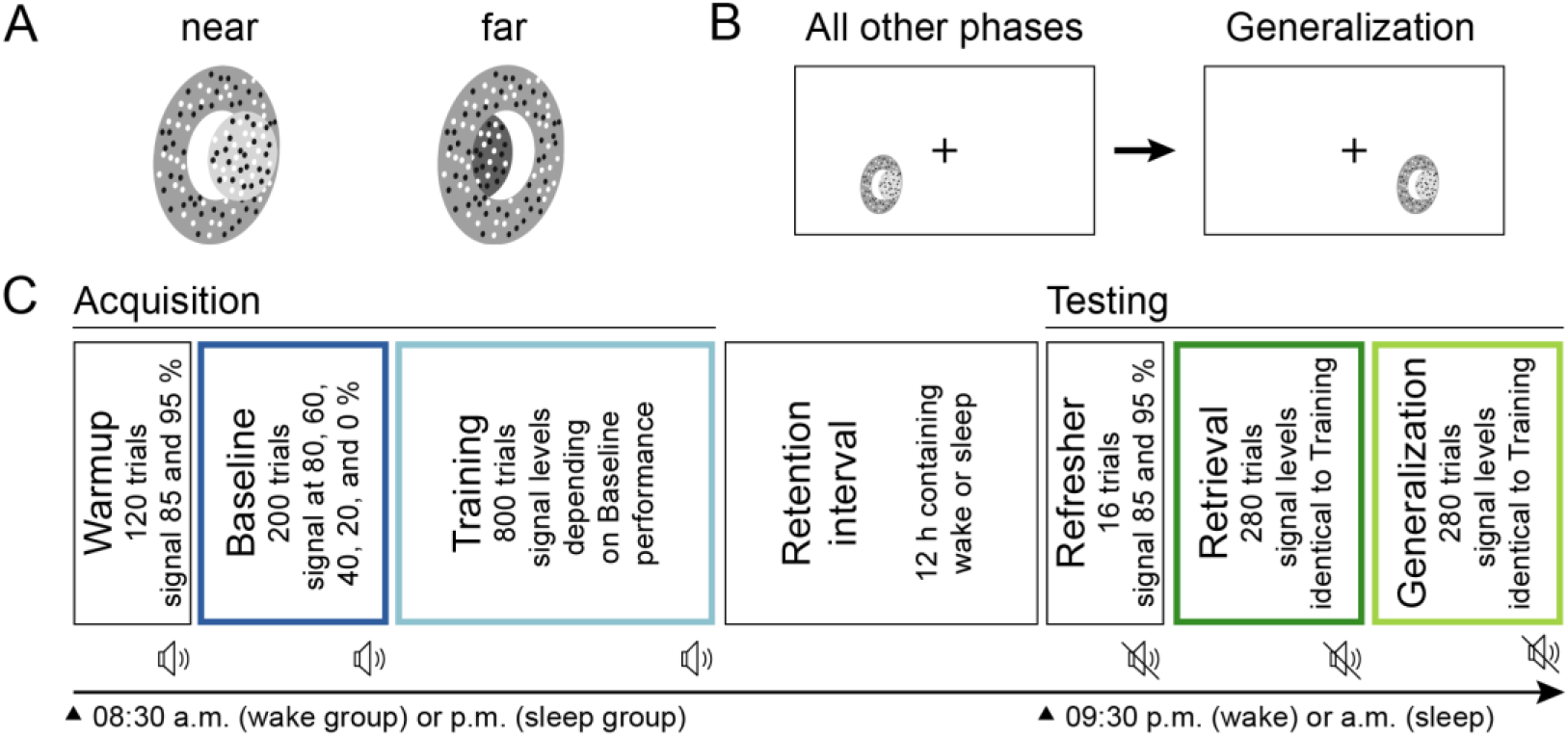
Learning task and experimental design. (A) Binocular disparity cues had to be exploited to differentiate two stimulus types (near vs. far). (B) All phases of the task were performed on one side of the lower visual field, except for the Generalization test, which was performed on the other side (balanced design). (C) Each participant went through an Acquisition and Testing session with a total of six experimental phases. Acquisition (Warmup, Baseline and Training) and Testing (Refresher, Retrieval and Generalization) were separated by a 13-h retention interval, resulting in a 12-h delay between Training and Retrieval. The wake group started Acquisition in the morning (8:30 a.m.), stayed awake during the retention interval, and was tested in the evening (09:30 p.m.). The sleep group started in the evening (08:30 p.m.), slept during retention, and completed the task in the morning (09:30 a.m.). The speaker symbol indicates whether tones signaled a correct and incorrect answer. This auditory feedback was given only during Acquisition. Phases used to assess performance are color-coded.

Disk and ring were assembled from circular dynamic random dot stereograms, consisting of equal numbers of black and white dots, the pattern of which was changed with each video-frame. Stimuli were shown on a back-projection screen using two projectors (Projection Design F21 DLP, 60 Hz, 1920×1080 pixels, 70.5 cm image width, 225 cd/m^2^ mean luminance, linearized, 80 cm viewing distance) and passive linear polarizing filters with a relative tilt of 90°. Participants wore passive linear polarizing filter glasses, allowing us to present slightly different images to each eye. The disparity between the two images resulted in the perception of depth, which we used to adjust the relative distance of the central disk to the outer ring, which in turn was always shown at 0° disparity. For each video frame (shown for 1/60 s), all dots belonging to the central disk had identical disparities. The disparity value for each frame was drawn randomly from a probability mass distribution set for the stimulus. Target disparities, determining the perceived distance between disk and ring (± 0.1° for “near” and “far”), were well above the detection threshold and thus easily detectable in the absence of disparity noise (noise disparity values were drawn from a uniform distribution, 11 values in 0.05° increments from −0.25° to 0.25°). Accordingly, task difficulty depended on the proportion of video frames (i.e., % signal strength) showing dots at the target disparity. For 0 % signal trials, disparity values were drawn only from the noise distribution and random feedback was given. We ran the task using custom scripts (cf. ref ^23^) using MATLAB 2014a (MathWorks, Natick, USA) and Psychophysics Toolbox 3^24,25^.

### Experimental timeline

All subjects participated in an Acquisition and a Testing session, which were scheduled 12 h apart and consisted of three phases each (Figure 1C). During Acquisition, we introduced participants to the task in a Warmup phase, during which stimuli were displayed with high signal strength (120 trials; signal at 85 or 95 %). During a Baseline phase, we linearly sampled the range of available signal strengths to assess the participant’s performance (200 trials; signal at 0, 20, 40, 60, and 80 %). Finally, in the Training phase, we presented stimuli with lower signal strengths, depending on their performance during Baseline. In addition to these rather difficult trials, we added one very high but rarely sampled signal strength to assess the subject’s lapse rate^26^ (800 trials with lower signal strengths being sampled more densely, e.g., 5, 10, 20, 40, and 90 %).

During Testing after the 12-hour retention period, we briefly refamiliarized subjects with the task (Refresher, 16 trials; signal at 85 and 95 %). Afterwards, Retrieval and Generalization were assessed with signal strengths identical to Training (280 trials each). During Generalization, the stimulus was presented again in the lower visual field but on the other side of the vertical meridian (i.e., moved from the left to the right side or vice versa, Figure 1 B). Trials were aborted in case of a fixation break during Warmup (after 40 initial trials that allowed free exploration), as well as during Baseline and the Refresher phase. During all other phases of the experiment, trials with fixation breaks were completed regularly but data from these trials were discarded from analysis (correct fixation rate in these phases: 91.34 +/− 6.95 %). Auditory feedback signaling correct vs. incorrect responses was only provided during Acquisition, it was disabled during Testing to minimize further learning.

Participants in the sleep group performed Acquisition in the evening, slept during the retention interval (7.48 ± 0.72 h) and went through Testing in the morning. Participants in the wake group started Acquisition in the morning, stayed awake, and completed Testing in the evening. We verified that subjects in the wake group did not nap during retention using activity trackers with acceleration and light sensors (Actiwatch, Philips Respironics, Murrysville, USA). The same devices were used to assess sleep duration in the sleep group. Any sleep in the wake group or sleep for less than 6 h in the sleep condition led to exclusion of the subject (n = 1). Subjects showed an increase in sleepiness across a wake retention interval (Holm-corrected p = 0.011, d = 3.469) and no change across a sleep retention interval (p = 1.0; interaction Acquisition/Testing × Sleep, p = 0.011, η^2^ = 0.079).

### Data analysis

We employed Bayesian inference (psignifit 4 toolbox for Matlab, https://github.com/wichmann-lab/psignifit) for estimating psychometric functions for Baseline, Training, Retrieval, and Generalization by fitting a beta-binomial model to the correct responses at each signal strength^27^. Based on the fitted cumulative Gaussian functions, we analyzed the slope of the psychometric function (assessed at the signal strength leading to 50 % correct responses), the threshold (signal strength required for 85 % performance, average for near/far), as well as bias (signal strength required for 50 % correct responses). On average, our models explained 96.55 % of the variance in the data (minimum: 84.70 %). Lapse rates were incorporated in the model and were generally very low (mean across subjects and experimental phases 2.17 ± 2.44 %).

Only trials in which participants correctly maintained fixation were analyzed. Outlier rejection was performed following an interquartile range outlier rejection rule with a multiplier of 2.2 (lower threshold: Q1 − 2.2 × (Q3 − Q1); upper threshold: Q3 + 2.2 × (Q3 − Q1))^28^. Statistics were performed using JASP 0.13 (https://jasp-stats.org) and by calculating repeated measures ANOVAs incorporating the within-subject factor Phase (Baseline/Training/Retrieval/Generalization) and the between-subject factor Wake/Sleep. In an additional analysis, we split the Training phase into three blocks of 260 trials each (Training 1-3, the first 20 trials were discarded). We confirmed results of the classical analysis by conducting additional Bayesian statistics. Since the first night of a previous experiment^19^ was identical to the current study, we aggregated data across studies for a second Bayesian analysis.

Post-hoc tests were corrected using Holm’s method. Results were Greenhouse-Geisser-corrected in case the sphericity assumption was violated. Effect sizes are provided for significant tests (η^2^ for ANOVA, Cohen’s d for post-hoc tests). Data were visualized using Python 3.7 and Seaborn 0.10.1. Error bars are shown as bootstrapped 95 % confidence intervals and, in case of skewed bootstrapped distributions, may be asymmetric. Error bars for Bayesian analyses reflect 95% credible intervals.

## Results

### Performance in the binocular disparity task improves locally

Task performance, as measured by the slope of the psychometric function, increased from Baseline to Retrieval (Figure 2, top). Performance data showed a trend towards steeper slopes from Baseline to Training (post-hoc paired-samples t-test, p = 0.152, Holm-corrected), a significant increase over the retention interval (Training vs. Retrieval, p = 0.006, d = 0.634), and a significant reduction when generalization of learning was tested in the opposite quadrant of the lower visual field (Retrieval vs. Generalization, p = 0.008, d = −0.596). As a result, performance during Generalization was not significantly different from Baseline levels, when participants were still naïve to the task (Generalization vs. Baseline, p = 0.268; ANOVA main effect Phase p < 0.001, η^2^ = 0.092). Interestingly, participants reached high performance levels early on during Training (Figure 2, bottom), resulting in no significant improvements across the three Training blocks (ANOVA main effect Training block, p = 0.567).

**Figure 2.**
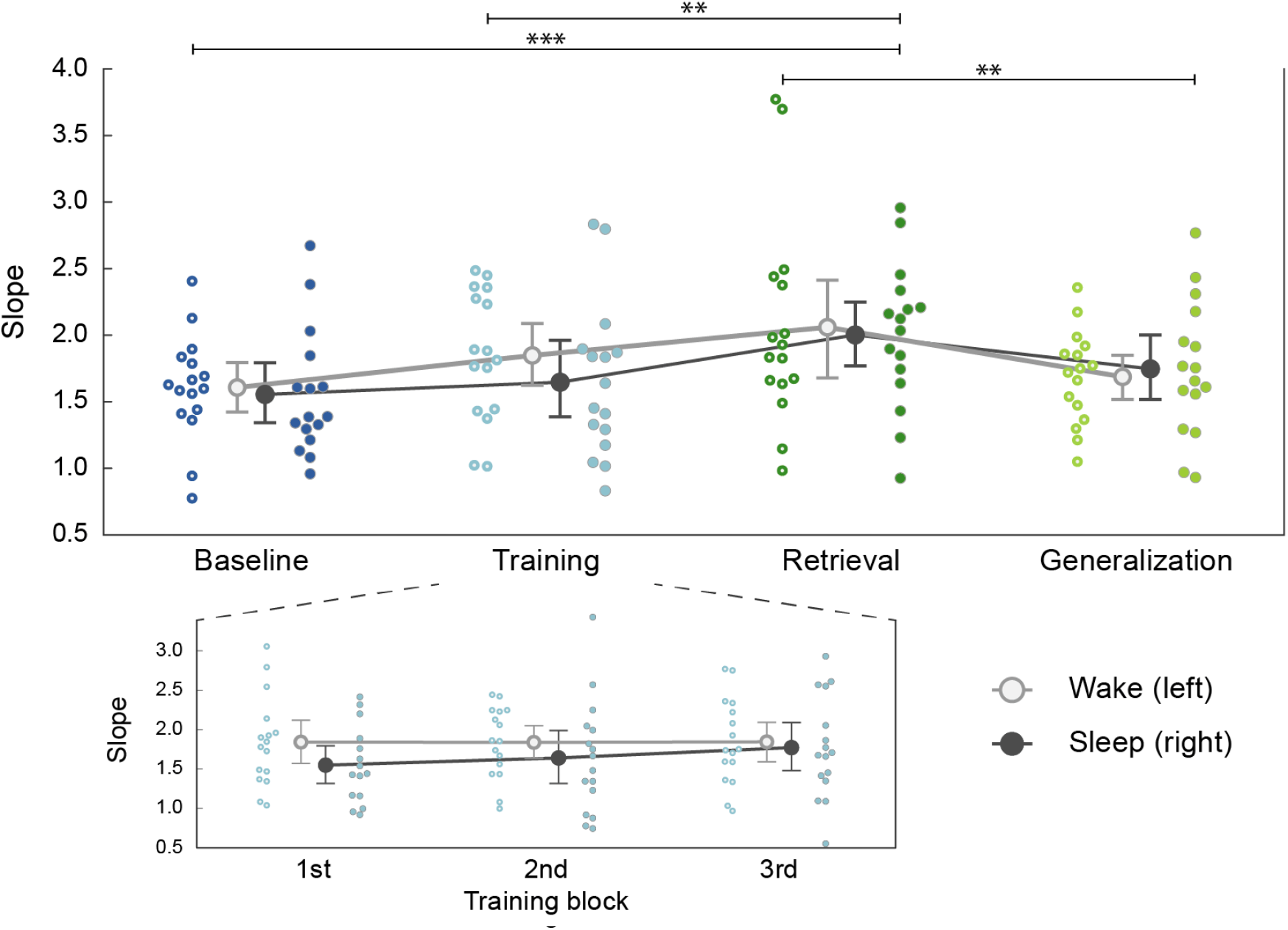
Improvements in binocular disparity do not generalize to the upper visual field. We compared the slope of the psychometric functions fitted to each participant’s performance between experimental phases and across groups (higher values denote an improvement in performance; for other parameters, see main text). Top: Large light unfilled and dark filled circles show grand averages (± 95 % confidence intervals) for the wake and sleep group, respectively. Small circles show single-subject data. Bottom: Training progress, illustrated by splitting Training trials into three consecutive blocks of 260 trials each. Performance improved only descriptively from Baseline to Training (post-hoc t-test, p = 0.152, Holm-corrected), improved significantly between Training and Retrieval (p = 0.006) and deteriorated when Generalization was tested in the upper visual field (p = 0.008; ANOVA main effect Phase p < 0.001). Performance did not differ between wake and sleep groups (ANOVA main effect Wake/Sleep, p = 0.688; interaction Wake/Sleep × Phase, p = 0.505) or across Training blocks (p = 0.567). ** p < 0.01; *** p < 0.001.

These findings were corroborated by results based on the psychometric threshold (ANOVA main effect Phase, p < 0.001, η^2^ = 0.091; performance improvements result in lower thresholds). Mirroring the slope data, threshold values significantly decreased over the retention interval (Baseline vs. Retrieval, p < 0.001, d = −0.907; Training vs. Retrieval, p = 0.006, d = −0.643) and increased again when the task was performed in the horizontally opposing visual-field quadrant (Retrieval vs. Generalization, p = 0.002, d = 0.740; all other p >= 0.332).

Finally, we analyzed bias, an observer’s inherent affinity towards one or the other response. As expected, participants’ bias changed over the course of the experiment (ANOVA main effect Phase: p = 0.009, η^2^ = 0.074), with the highest values observed during Baseline, when subjects were still unaccustomed to the task, and subsequent decreases for Training and Retrieval (Baseline vs. Training, p = 0.016, d = −0.577; Baseline vs. Retrieval, p = 0.070, d = −0.444). Bias increased slightly during Generalization, but this change was significant only when compared to Training, which had the lowest bias values (Training vs. Generalization, p = 0.018, d = 0.556; all other p >= 0.29).

### No signs for improved generalization after sleep

So far, we have demonstrated that changes in performance across the retention period did not differ between sleep and wake groups at the trained location. We now analyzed whether sleep increased the participants’ ability to generalize their skill improvements to the other visual hemifield. In contrast to our previous study^19^, we tested generalization across the vertical instead of the horizontal meridian. This is a decisive difference because anisotropies between the lower and upper visual field may have confounded our earlier results. In line with our previous findings, neither performance nor bias evolved differently across the experiment in the wake and sleep groups (ANOVA main effect Wake/Sleep: Slope, p = 0.688; Threshold, p = 0.330; Bias, p = 0.996; interaction Wake/Sleep × Phase: Slope, p = 0.505; Threshold, p = 0.198; Bias, p = 0.626). Similarly, a planned direct comparison of performance and bias in the sleep and wake groups during Generalization showed no significant differences (all uncorrected p > 0.542).

To substantiate the notion that sleep does not influence training and generalization success, we performed Bayesian ANOVAs on all parameters. As Bayesian inference is based on model comparison, this allows us to quantify the empirical evidence in favor of the null hypothesis, something that is conceptually impossible using classical statistics Analyses were conducted using JASP’s default priors (r scale = 0.5 for fixed effects and 1.0 for random effects). Given the highly significant main effects of Phase, we included this factor into the respective null models and compared them to full models that additionally contained the main effect of Wake/Sleep and its interaction with Phase. These analyses provided moderate to strong evidence in favor of the null models (slope, BF_01_ = 13.15; threshold, BF_01_ = 4.11; bias, BF_01_ = 23.10). Subsequent analyses of individual effects indicated mostly anecdotal evidence against a main effect of Wake/Sleep (slope, BF_excl_ = 2.35; threshold, BF_excl_ = 1.83; bias, BF_excl_ = 3.31), but more reliable evidence against the interaction of Wake/Sleep and Phase (slope, BF_excl_ = 5.46; threshold, BF_excl_ = 2.23; bias, BF_excl_ = 7.05). This pattern of results was even more pronounced when the first session of all 17 participants from our previous study^19^ were included in the analysis (slope, BF_01_ = 19.08; threshold, BF_01_ = 11.73; bias, BF_01_= 28.90), with moderate evidence against both the main effect of Wake/Sleep (slope, BF_excl_ = 3.15; threshold, BF_excl_ = 2.94; bias, BF_excl_ = 3.45) and its interaction with Phase (slope, BF_excl_ = 6.05; threshold, BF_excl_ = 3.99; bias, BF_excl_ = 8.37).

Both classical and Bayesian statistics are thus compatible with a lack of an important effect of sleep on the generalization of improvements in coarse disparity discrimination. This result has now proved consistent across two independent sets of participants (total n = 49).

## Discussion

The data reported here confirm our previous findings based on the same stimuli and task^19^. They show substantial improvements in coarse binocular disparity discrimination following a single training session. Our results also corroborate that these learning effects are retinotopically local, i.e. they do not transfer between different visual hemifields. In our previous study, we investigated transfer from the lower to the upper hemifield^19^. The lack of generalization observed there could have been partly the result of anatomic and functional anisotropies between the upper and lower visual hemifields^21,22,29,30^. Here, we addressed this potential confound by assessing transfer between horizontal hemifields and observe qualitatively and quantitatively similar effects. In particular, when controlling for low-level differences between training and test locations in this way, we again find that sleep during the retention interval does not affect subsequent performance, neither within the trained visual-field quadrant nor at the transfer location.

These results suggest that perceptual learning of coarse disparity discrimination depends on retinotopically specific mechanisms that do not benefit from memory consolidation during subsequent sleep. While performance increased significantly over the retention period, this increase did not differ between sleep or wakefulness and improvements remained local in both experimental groups. Thus, disparity discrimination learning does not seem to be amenable to transfer via either sleep-dependent extraction of higher-level regularities^31^, improved temporal allocation of attention^17^, or other sleep-dependent top-down processes that improve task performance. In this context, it seems unlikely that improved read-out of noisy low-level signals and subsequent optimization of decision criteria at higher processing levels can account for the present results^32^. Optimization at higher levels would be expected to result in less local specificity after either sleep or wakefulness than seen in this study.

As discussed previously, both retinotopic specificity and independence of sleep may be explained by the nature of our stimuli^19^: since difficulty was manipulated in terms of the number of frames containing the target disparity, optimizing performance presumably involved a combination of improving the tuning of neurons coding for the target disparity and reducing interference by non-informative noise frames^33^. In this context, integrating signal and discarding noise over the 1.5 s trial duration may depend on early sensory buffers^34^ that do not benefit from sleep-dependent reactivation of neuronal ensembles. Results may differ in paradigms in which the temporal structure within trials supports task performance or in which the task involves stimuli that are either static or change on a time scale comparable to natural scenes.

Interestingly, we find that behavioral performance improved rapidly and plateaued early on during training. Indeed, psychophysical indicators of performance remained statistically unchanged throughout the training session, and the significant increase in performance from baseline to retrieval was mostly the result of improvements that occurred during the retention interval. In combination with the absence of sleep effects, this suggests that the passage of time alone is sufficient for binocular disparity discrimination learning to be consolidated. However, adaptation and fatigue are frequently reported to impair performance during extended perceptual training sessions^35,36^. Such effects may have masked learning during the training session, so that performance towards the end of the training session was underestimated. Performance increases over the retention interval may thus reflect a combination of a release from adaptation and fatigue, as well as genuine time-dependent improvements. Nevertheless, our results suggest substantial enhancements in disparity discrimination across a 12-h retention interval without further practice, independent of participants’ behavioral state during this interval.

The lack of any kind of interaction between consolidation and sleep may appear surprising given a long history of perceptual learning studies which have documented beneficial effects of sleep^6–13^. However, practically all demonstrations of sleep effects were obtained with a single experimental paradigm, the TDT originally developed by Karni and Sagi^4^. As outlined previously^19^, the nature of this particular task differs substantially from our protocol, as well as from many other tasks classically used to investigate perceptual learning^37,38^. Most importantly, difficulty in the classical TDT is manipulated, and learning is induced, by reducing the SOA between target and mask displays. Recent evidence suggests that learning on the TDT largely relates to improved temporal learning^17^ and that appropriate training procedures can induce transfer of texture discrimination learning. This could explain why a growing number of studies report substantial amounts of ocular and retinotopic transfer^31,39,40^, in contrast to findings based on the classical paradigm^4,41^. A promising avenue for future research would be to use a more strictly low-level task such as the present one and to manipulate individual stimulus or task dimensions to probe the boundary conditions for learning to become transferable and/or sleep-dependent. For example, it would be interesting to test whether the dynamic nature of our stimuli prevents additional improvement and/or transfer via sleep-dependent reactivation, or whether selectively rewarding performance for particular stimulus classes would lead to selective and sleep-dependent learning^42^.

The main reason for conducting this follow-up study was to account for the potentially confounding effects of numerous visual-field anisotropies^30,43,44^, particularly those between the vertical hemifields^21,22,29,45,46^. For example, differences in spatial and temporal resolution between upper and lower visual fields could have masked high-level transfer of learning, as well as sleep benefits, given that participants were always trained in the lower visual field. Furthermore, some retinotopic specializations are directly linked to disparity processing, such as preferences for crossed vs. uncrossed disparities in the lower vs. upper visual field^21^. The present data suggest that these anisotropies do not affect binocular disparity discrimination learning. While there are a number of horizontal asymmetries in visual processing^47–49^, these appear much less likely to exert a substantial influence in the context of our stimuli and task^50,51^. Thus, we are confident that the present data, in combination with our previous results^19^, provide strong evidence for retinotopically local learning of disparity discrimination that is independent of both sleep and the particular combination of training and testing locations.

While binocular disparity discrimination is well-suited for our particular research question, it is unclear how far the present results can be generalized to other tasks, such as contrast detection, motion detection, or fine form judgements^52^. More generally, future studies should test for sleep effects on a wider range of perceptual skills, in order to clarify whether sleep benefits reported for the TDT are the rule or an exception. Another limitation of the present study results from mirroring trade-offs made for our previous study^19^. Here, we tested sleep effects in a between-subject design. This allowed us to perform both Training and Testing in the lower visual field without occupying more than one location per visual quadrant, ruling out visual field anisotropies as a potential confounding factor. In the previous study, subjects performed the task in all four quadrants, allowing for a statistically more powerful within-subject design.

Research on perceptual learning has made fundamental contributions to our understanding of how the nervous system balances the competing needs for plasticity and stability^53–55^. The overall picture emerging from literally thousands of studies is of a surprising degree of flexibility in low-level sensory processing, even in the adult brain^56^. However, the boundary conditions for this flexibility are much more restricted than for higher forms of learning^57,58^. Recently, Gilead et al. (2020)^59^ proposed that a better understanding of mind and behavior may require more detailed investigations into how neural systems abstract information. In their framework, perceptual learning as investigated here constitutes the simplest case of *modality-specific abstraction* across stimulus features, individual objects, and their relations. Against this background, it is tempting to speculate that both sleep-dependent memory consolidation and substantial transfer of learning may occur only when higher-level, at least multimodal, abstraction is required for successful performance. Importantly, such relatively simple instances of abstraction are likely those with the longest evolutionary history^60^ and may thus provide a solid basis for comparative research into perception and learning. Delimiting the conditions necessary and sufficient for low-level learning to be re-processed and generalized would be a crucial step towards integrative models of robust perception^61^.

## Acknowledgments

This research was supported by grants from the German Research Foundation (Deutsche Forschungsgemeinschaft, DFG): SFB 1233 (project no. 276693517) and a Research Fellowship (project no. 430174808).

## Additional Information

The authors declare no competing interests.

## Author contributions statements

JGK and KR designed the study. HN provided the experimental setup and code. JGK coordinated the study, carried out the experiments, and analyzed the data. JGK and KR carried out the statistical analyses and drafted the manuscript. All authors reviewed the manuscript and gave final approval for publication.

